# A resource of targeted mutant mouse lines for 5,061 genes

**DOI:** 10.1101/844092

**Authors:** Marie-Christine Birling, Atsushi Yoshiki, David J Adams, Shinya Ayabe, Arthur L Beaudet, Joanna Bottomley, Allan Bradley, Steve DM Brown, Antje Bürger, Wendy Bushell, Francesco Chiani, Hsian-Jean Genie Chin, Skevoulla Christou, Gemma F Codner, Francesco J DeMayo, Mary E Dickinson, Brendan Doe, Leah Rae Donahue, Martin D Fray, Alessia Gambadoro, Xiang Gao, Marina Gertsenstein, Alba Gomez-Segura, Leslie O Goodwin, Jason D Heaney, Yann Hérault, Martin Hrabe de Angelis, Si-Tse Jiang, Monica J Justice, Petr Kasparek, Ruairidh E King, Ralf Kühn, Ho Lee, Young Jae Lee, Zhiwei Liu, K C Kent Lloyd, Isabel Lorenzo, Ann-Marie Mallon, Colin McKerlie, Terrence F Meehan, Stuart Newman, Lauryl MJ Nutter, Goo Taeg Oh, Guillaume Pavlovic, Ramiro Ramirez-Solis, Barry Rosen, Edward J Ryder, Luis A Santos, Joel Schick, John R Seavitt, Radislav Sedlacek, Claudia Seisenberger, Je Kyung Seong, William C Skarnes, Tania Sorg, Karen P Steel, Masaru Tamura, Glauco P Tocchini-Valentini, Chi-Kuang Leo Wang, Hannah Wardle-Jones, Marie Wattenhofer-Donzé, Sara Wells, Brandon J Willis, Joshua A Wood, Wolfgang Wurst, Ying Xu, IMPC Consortium, Lydia Teboul, Stephen A Murray

## Abstract

The International Mouse Phenotyping Consortium reports the generation of new mouse mutant strains for over 5,000 genes from targeted embryonic stem cells on the C57BL/6N genetic background. This includes 2,850 null alleles for which no equivalent mutant mouse line exists, 2,987 novel conditional-ready alleles, and 4,433 novel reporter alleles. This nearly triples the number of genes with reporter alleles and almost doubles the number of conditional alleles available to the scientific community. When combined with more than 30 years of community effort, the total mutant allele mouse resource covers more than half of the genome. The extensively validated collection is archived and distributed through public repositories, facilitating availability to the worldwide biomedical research community, and expanding our understanding of gene function and human disease.

## Results and Discussion

Despite thirty years of mouse targeted mutagenesis, *in vivo* function of the majority of genes in the mouse genome are still unknown. This reflects the observation that a small number of genes have been the object of intensive study including the development of multiple mouse models, while a significant proportion of the coding genome remains entirely unexplored ^1^. The completion of the sequencing of the mouse genome, coupled with the use of mouse embryonic stem (ES) cells for gene targeting to create complex mutant alleles, presented an opportunity to functionally analyze all the protein coding genes of a mammalian species ^2,3^. Taking advantage of comprehensive manual annotation of the genome ^4^, the International Knockout Mouse Consortium (IKMC) systematically generated single-gene, reporter-tagged null alleles for protein-coding genes by homologous recombination in mouse ES cells ^5,6^. Subsequently, large-scale mouse production and phenotyping programs deployed these unique resources, establishing the feasibility of genome-scale mouse production and phenotyping ^7–9^. Building upon these successes, the International Mouse Phenotyping Consortium (IMPC) was established to coordinate a network of programs around the globe, assuring uniformity and reproducibility of these efforts, including standardization of phenotyping protocols and the use of a single inbred mouse strain background, C57BL/6N, with the ultimate goal of generating and phenotyping a single-gene knockout (KO) mouse line for every protein-coding gene in the genome.

Production of KO mice began in concert with the expansion of the ES cell library, but rapidly accelerated after 2011 with the funding of multiple IMPC programs. To date, more than 17,500 individual production attempts (microinjection or aggregation) have resulted in the germline transmission of KO alleles for 5,061 unique genes (**Figure 1A**; **Supplemental Table ST1**). These lines have been expanded for phenotyping, providing key insights into mammalian biology and disease ^10–15^; www.mousephenotype.org). The IMPC contribution extends the total number of genes with targeted KO alleles produced by the scientific community from the 8,391 reported and curated by Mouse Genome Informatics (MGI; ^16^)(**Figure 1B**; **Supplemental Table ST2**), to 11,241, or more than half of the genome. Much of the overlap (2,211 genes) reflects specific community requests for the production of novel complex alleles (see below), targeting on an inbred C57BL/6N background, or for mutant mouse lines unavailable through public repositories. The growing use of CRISPR/Cas9 editing to produce null alleles for the IMPC led to the decrease in ES cell-based production beginning in 2015.

**Figure 1:**
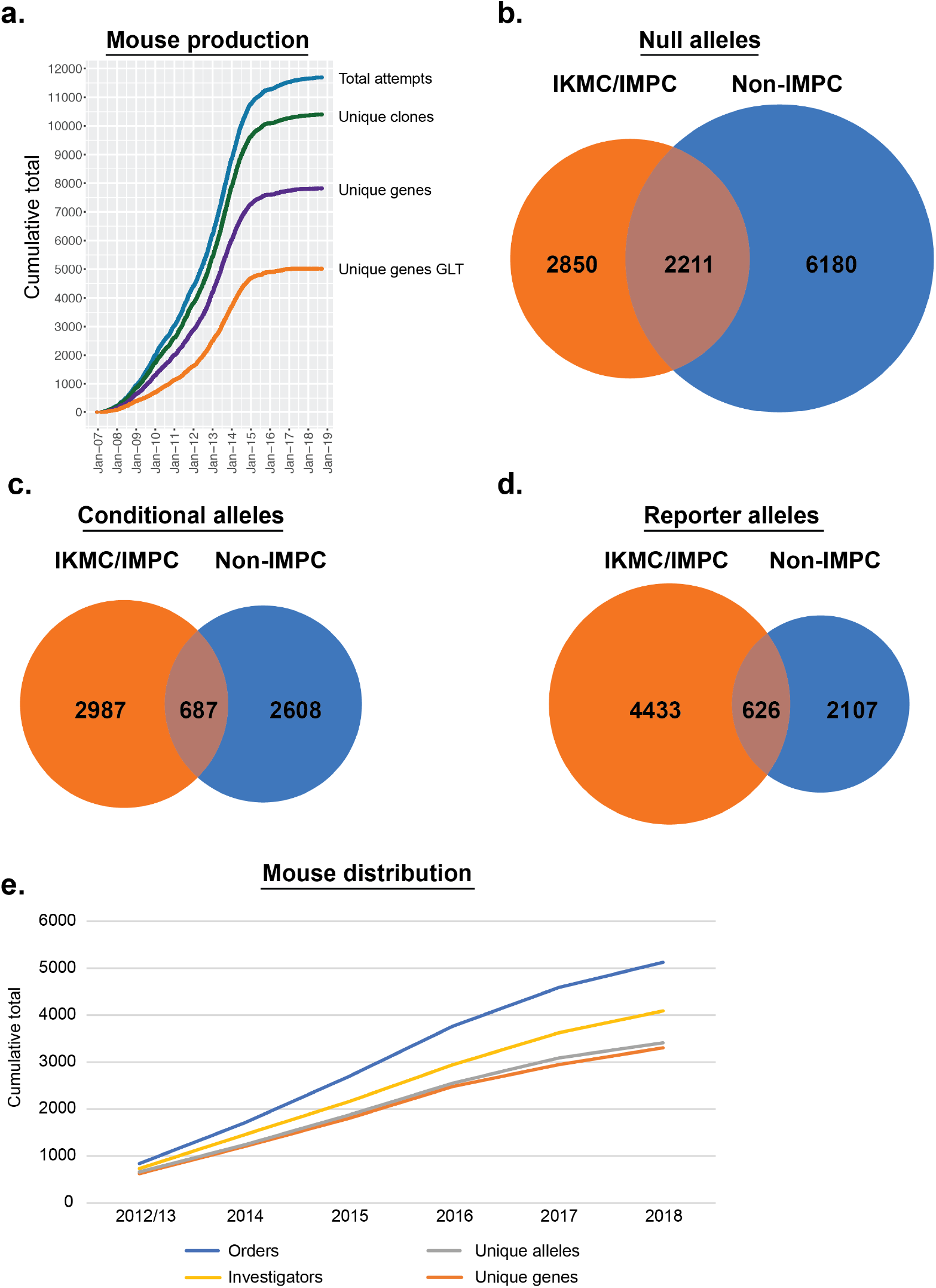
Generation and impact of targeted alleles for 5,061 unique mouse genes. (a) Cumulative production progress, including all attempts (microinjection or aggregation (black), unique ES cell clones injected (red), unique genes attempted (yellow), and unique genes that achieved germline transmission (GLT; blue). For GLT, the date reflects the date of microinjection, and only reports the first instance of transmission for the small number of duplicate mutations produced. (b) Venn representation of unique gene null alleles produced by the IMPC (orange) and by the rest of the scientific community as reported in MGI (“Non-IMPC”; blue). (c) Unique gene conditional-ready alleles produced by the IMPC (orange) and by the rest of the scientific community (blue). (d) Unique gene reporter alleles produced by the IMPC (orange) and by the rest of the scientific community (blue). (e) Cumulative mouse orders of IMPC lines processed by production centres and mouse model Repositories from 2012-2018 (blue line). The cumulative number of ordering investigators, unique alleles ordered, and unique genes ordered are shown in yellow, grey, and orange, respectively.

While the primary goal of the IKMC and IMPC was to generate and phenotype a null allele for every protein-coding gene, the mutant alleles included additional functional features. All alleles included a *lacZ* reporter cassette to facilitate analysis of gene transcription *in situ* (**Supplemental Figure S1**; ^5,6^). A large proportion of the alleles have conditional potential, providing future users with a useful tool for detailed, mechanistic analyses (Supplemental Figure S1a). The multifunctional utility of the alleles produced by the IMPC has greatly expanded the repertoire of genetic resources available to the scientific community. Of the 3,674 unique gene, conditional-ready mouse models generated and validated, 2,987 were novel alleles for genes without an existing conditional allele (81.3%). These nearly double the total number of genes with conditional KO alleles produced by the scientific community as a whole (2,987 IMPC conditional alleles added to the 3,295 conditional alleles reported in MGI as mouse lines; **Figure 1C**). The impact is even more significant for reporter alleles. The IMPC has produced reporter alleles for 5,059 unique genes, of which 4,433 are novel (87.6%), complementing the 2,733 produced by the scientific community (**Figure 1D**). This has nearly tripled the total number of genes with reporter alleles available to the community as mouse lines.

The generation of mouse lines was underpinned by comprehensive quality control strategies for both ES cell karyotype and targeted allele, which ensured efficient production and integrity of the targeting event (**Supplemental Table ST1** and **ST2**; **Supplemental Figures S2** and **S3**). Further quality control (QC) analysis also revealed that part of the ES cell collection contained an additional insertion of a wild-type *nonagouti* (*A*) gene on chromosome 8, likely introduced with the targeted reversion event in these cell lines (subclone JM8A3^17^). However, as the insertion of the wild-type *nonagouti* gene results in an agouti coat color, this allele can be easily segregated from the mutant allele in most cases (**Supplemental Figure S4**). High-throughput allele validation of ES cells was performed using either a suite of quantitative and endpoint PCR-based tests or a combination of Southern blot and PCR-based analysis, depending on production centre (**Supplemental Figure S2, S3** and **S5** and **Supplemental Tables ST3** and **ST4** and Methods). Despite these efforts, we found that additional quality control (QC) on the mouse lines themselves was required to ensure all IMPC lines resulted from the transmission of the correctly targeted allele (**Supplemental Table ST5**; ^18^). This additional QC at the mouse level identified a small but significant proportion of incorrect alleles that transmitted through the germline of chimera mice derived from clones that had passed initial and secondary validation QC testing in the ES cell. Our experience highlights the importance of careful allele validation before and after mouse production.

As a result of this effort, mouse lines with targeted alleles for more than 5,000 genes on a C57BL/6N genetic background with extensive and documented genetic validation of the targeted locus are now available to the biomedical research community, supporting high standards of reproducibility for future investigations. The IMPC resource has shown its usefulness through the continued and robust uptake of mutant mouse lines by investigators around the world. This includes both KO and conditional alleles with mouse lines distributed as live mice and cryopreserved stocks. To date, over 5,000 orders for mutant mice for 3,301 unique genes have been processed and shipped to more than 4,000 investigators around the world (**Figure 1E**). To date, more than 1,900 publications acknowledge the use of EUCOMM/KOMP alleles (for example ^19–30^). This demonstrates the utility of these resources, the cumulative use of which continues to grow over time, and complements the systematic phenotyping efforts of IMPC centres. In the new era of genome editing, this ES cell-derived collection remains of unique value as it offers particularly sophisticated and quality-controlled alleles representing a cornerstone of the collective development of a null allele resource for the complete mammalian genome ^2^.

### Data availability

All data are freely available from the IMPC database hosted at EMBL-EBI via a web portal (mousephenotype.org), ftp (ftp://ftp.ebi.ac.uk/pub/databases/impc) and automatic programmatic interfaces. An archived version of the database will be maintained after cessation of funding (exp. 2021) for an additional 5 years. Information on alleles, together with phenotype summaries, are additionally archived with Mouse Genome Informatics at the Jackson Laboratory via direct data submissions (J:136110, J:148605, J:157064, J:157065, J:188991, J:211773).

## Acknowledgments

We thank all technical personnel at the different centres involved in this project for their contribution. We thank Gareth Clarke for support with illustrations. M.-C.B., G.P., M. W.-D., Y.H., T.S., were supported by the Université de Strasbourg, the CNRS, the INSERM and the programmes ‘Investissements d’avenir’ (ANR-10-IDEX-0002-02, ANR-10-LABX-0030-INRT, ANR-10-INBS-07 PHENOMIN), A.Y. and M.T. were supported by RIKEN BioResource Research Center (BRC), Management Expenses Grant from the Ministry of Education, Culture, Sports, Science and Technology (MEXT), D.A, J.B., A.B., W.B, B.D., S.N, R.R.S., B.R., E.R., W.S., K.S. and H.W.-J. were supported by the Wellcome Trust, S.B., S.C., G. C., M.F., S.W. and L.T. were supported by the Medical Research Council, C.McK. and L.N. were supported by Genome Canada and Ontario Genomics (OGI-051),), H.L., Y.J.L., G.T.O., J.K.S. supported by National Research Foundation (2014M3A9D5A01074636, 2014M3A9D5A01075128), Republic of Korea (KMPC), M.B., A. B., S.B., A.B., W.B., F.C., M.F., A.G., M. H. A., R. K., S.N., G.P., R.R.S., B.R., E.R., J.S., W.S., C.S., T.S., G.T.-V, S.W., W.W. and L.T. were supported by the European Commission (EUCOMM, EUMODIC, Infrafrontier). R.S. and P.K. were supported by RVO 68378050 by Academy of Sciences of the Czech Republic and by LM2015040 (Czech Centre for Phenogenomics), CZ.1.05/2.1.00/19.0395, CZ.1.05/1.1.00/02.0109 funded by the Ministry of Education, Youth and Sports and the European Regional Development Fund. Work was supported by the National Key R&D Program of China 2018YFA0801100 to Y.X. Research reported in this publication was supported by the NIH Common Fund, the Office of The Director, and the National Human Genomic Research Institute of the National Institutes of Health under Award Numbers (U42OD011174 supported J.B., A.B., W.B., B.D., M.D., M.F., J. H., M. J., I.L., F.M., S.N, R.R.S. B.R. E.R. W.S., J.S., S.W.; U42OD011175 supported M.G., C.McK., L.N., B.W., J.W, K.C.L.; U42OD011185 supported L.R.D., L. G. and S.A.M.; U54HG006370-02 supported A. G-S., R. K., A.-M.M., T.M., L.S.). The content is solely the responsibility of the authors and does not necessarily represent the official views of the National Institutes of Health.

## Author Contributions

M-C.B., A.Y., S.A., J.B., A.Bu., W.B., F.C., S.C., G.F.C., F.J.D., B.D., M.D.F., A.G., M.G., A.G-S., L.O.G., R.E.K., R.K., H.L., Y.J.L., I.L., A-M. M., C.M., T.F.M., S.N., L.M.J.N., G.T.O., G.P., R.R-S., B.R., E.J.R., L.A.S., J.S., J.R.S., C.S., H.W-J., M.W-D., B.J.W., J.A.W., L.T., and S.A.M generated data, developed data tools and databases, and/or carried out data and statistical analyses; A.Y., D.J.A., A.L.B, A.B., S.D.M.B., H-J.G.C., M.E.D., L.R.D., X.G., J.D.H., Y.H., M.H.A., S-T.J., M.J.J., Z.L., K.C.K.L., J.K.S., W.C.S., T.S., K.P.S., M.T., G.P.T-V., C-K.L.W., S.W., W.W., Y.X., L.T., and S.A.M. directed research at their respective institutions; M-C B., L.T., and S.A.M wrote the paper.

## Additional Information

Supplementary Methods, Figures, and Tables accompany this manuscript.

## References

1. Stoeger, T., Gerlach, M., Morimoto, R.I. & Nunes Amaral, L.A. Large-scale investigation of the reasons why potentially important genes are ignored. PLoS Biol 16, e2006643 (2018).

2. Austin, C.P. et al. The knockout mouse project. Nat Genet 36, 921–4 (2004).

3. Auwerx, J. et al. The European dimension for the mouse genome mutagenesis program. Nat Genet 36, 925–7 (2004).

4. Ashurst, J.L. et al. The Vertebrate Genome Annotation (Vega) database. Nucleic Acids Res 33, D459–65 (2005).

5. Skarnes, W.C. et al. A conditional knockout resource for the genome-wide study of mouse gene function. Nature 474, 337–42 (2011).

6. Valenzuela, D.M. et al. High-throughput engineering of the mouse genome coupled with high-resolution expression analysis. Nat Biotechnol 21, 652–9 (2003).

7. de Angelis, M.H. et al. Analysis of mammalian gene function through broad-based phenotypic screens across a consortium of mouse clinics. Nat Genet 47, 969–78 (2015).

8. White, J.K. et al. Genome-wide generation and systematic phenotyping of knockout mice reveals new roles for many genes. Cell 154, 452–64 (2013).

9. West, D.B. et al. A lacZ reporter gene expression atlas for 313 adult KOMP mutant mouse lines. Genome Res 25, 598–607 (2015).

10. Bowl, M.R. et al. A large scale hearing loss screen reveals an extensive unexplored genetic landscape for auditory dysfunction. Nat Commun 8, 886 (2017).

11. Dickinson, M.E. et al. High-throughput discovery of novel developmental phenotypes. Nature 537, 508–514 (2016).

12. Karp, N.A. et al. Prevalence of sexual dimorphism in mammalian phenotypic traits. Nat Commun 8, 15475 (2017).

13. Meehan, T.F. et al. Disease model discovery from 3,328 gene knockouts by The International Mouse Phenotyping Consortium. Nat Genet 49, 1231–1238 (2017).

14. Munoz-Fuentes, V. et al. The International Mouse Phenotyping Consortium (IMPC): a functional catalogue of the mammalian genome that informs conservation. Conserv Genet 19, 995–1005 (2018).

15. Rozman, J. et al. Identification of genetic elements in metabolism by high-throughput mouse phenotyping. Nat Commun 9, 288 (2018).

16. Bult, C.J. et al. Mouse Genome Database (MGD) 2019. Nucleic Acids Res 47, D801–D806 (2019).

17. Pettitt, S.J. et al. Agouti C57BL/6N embryonic stem cells for mouse genetic resources. Nat Methods 6, 493–5 (2009).

18. Ryder, E. et al. Molecular characterization of mutant mouse strains generated from the EUCOMM/KOMP-CSD ES cell resource. Mamm Genome 24, 286–94 (2013).

19. Ingham, N.J. et al. Mouse screen reveals multiple new genes underlying mouse and human hearing loss. PLoS Biol 17, e3000194 (2019).

20. Liu, X. et al. The complex genetics of hypoplastic left heart syndrome. Nat Genet 49, 1152–1159 (2017).

21. Miyata, H. et al. Genome engineering uncovers 54 evolutionarily conserved and testis-enriched genes that are not required for male fertility in mice. Proc Natl Acad Sci U S A 113, 7704–10 (2016).

22. Pol, A. et al. Mutations in SELENBP1, encoding a novel human methanethiol oxidase, cause extraoral halitosis. Nat Genet 50, 120–129 (2018).

23. Small, K.S. et al. Regulatory variants at KLF14 influence type 2 diabetes risk via a female-specific effect on adipocyte size and body composition. Nat Genet 50, 572–580 (2018).

24. Akawi, N. et al. Discovery of four recessive developmental disorders using probabilistic genotype and phenotype matching among 4,125 families. Nat Genet 47, 1363–9 (2015).

25. Bertero, A. et al. Activin/nodal signaling and NANOG orchestrate human embryonic stem cell fate decisions by controlling the H3K4me3 chromatin mark. Genes Dev 29, 702–17 (2015).

26. de la Rosa, J. et al. A single-copy Sleeping Beauty transposon mutagenesis screen identifies new PTEN-cooperating tumor suppressor genes. Nat Genet 49, 730–741 (2017).

27. Kim, J.H. et al. Regulation of the catabolic cascade in osteoarthritis by the zinc-ZIP8-MTF1 axis. Cell 156, 730–43 (2014).

28. Koch, S., Acebron, S.P., Herbst, J., Hatiboglu, G. & Niehrs, C. Post-transcriptional Wnt Signaling Governs Epididymal Sperm Maturation. Cell 163, 1225–1236 (2015).

29. Kochubey, O., Babai, N. & Schneggenburger, R. A Synaptotagmin Isoform Switch during the Development of an Identified CNS Synapse. Neuron 90, 984–99 (2016).

30. Lin, C.J., Koh, F.M., Wong, P., Conti, M. & Ramalho-Santos, M. Hira-mediated H3.3 incorporation is required for DNA replication and ribosomal RNA transcription in the mouse zygote. Dev Cell 30, 268–79 (2014).

