## Supplemental Material for "A resource of targeted mutant mouse lines for 5,061 genes"

**Supplemental Materials**

Supplemental Tables ST1 and ST2 are provided as separate Excel files

Page 2-3: Supplemental Methods

Page 4-5: Supplemental Figure S1

Page 6: Supplemental Figure S2

Page 7: Supplemental Figure S3

Page 8: Supplemental Figure S4

Page 9: Supplemental Figure S5

Page 10: Supplemental Table ST3

Page 11: Supplemental Table ST4

Page 12: Supplemental References

Page 13: Animal Welfare Assurances

Page 14-15: IMPC Contributors

**Supplemental Methods**

**Data analysis**

All IMPC production data was downloaded from the International Micro-Injection Tracking System database (iMITS) (www.mousephenotype.org/imits) and manually curated to remove unrelated projects. Attempts were further cross-referenced with Mouse Genome Informatics (MGI) to assure all alleles were publicly reported. For the small number of cases where multiple alleles and/or mouse lines were generated for the same gene, the first successful attempt was selected, resulting in a final list of 5,061 genotype confirmed mutant genes (**Supplementary Table 1**). For lists of mice with targeted mutations produced by the scientific community, MouseMine (www.mousemine.org) was used to query the Mouse Genome Database [1] (MGD) to identify all targeted, endonuclease-mediated, ENU, gene-trapped, and transposon-induced alleles that were either null, potentially null (conditional), or reporter null. A full list of allele attributes as well as a list of all null, conditional, and reporter alleles, is provided in **Supplemental Table 2**. Mice produced using ES cells from IKMC resources that overlapped iMITS data were excluded, while all other lines produced using these clones were included in the community list.

**Mouse availability**

All mouse lines established and validated by the International Mouse Phenotyping Consortium are available for distribution through public repositories as indicated on the IMPC portal (www.mousephenotype.org).

**Allele validation**

PCR-based validation methods were previously described for Velocigene (Vlcg) [2] and KO-first allele types [3]. For Southern blot-based QC, allele validation of ES cells was performed on phenol/chloroform extracted genomic DNA. Two restriction enzymes for each side of the locus were chosen outside of each homology arm to produce DNA segments encompassing the whole of each homology arm of the targeting construct and either the *lacZ* or the *neo* cassette. Southern blot hybridization was performed with corresponding *lacZ* or *neo* probes, radioactively- or DIG-labelled. A unique band of a given size was expected with each enzyme. The pattern formed by the four digests yielded a profile specific for a given targeting event. Bands of unexpected size or multiple bands for at least two enzymes constituted a failure.

**ES cell to mouse conversion**

Whenever operationally possible, the chromosome count of ES cells was assayed by either chromosome spreads or quantitative PCR (qPCR or ddPCR) prior to microinjection or aggregation [4] either at the ES cell Repository (www.komp.org; www.eummcr.org) or at individual production centres. ES cells were cultured either in standard ES cell media [5] or in media with serum replacement and two inhibitors, mitogen-activated protein kinase inhibitor PD0325901 and glycogen synthase kinase-3 inhibitor CHIR99021 (2i media; [6]) prior to chimera generation. Chimera generation was performed by either microinjection of ES cells into blastocyst or morula hosts or ES cell-morula aggregation [5,6]. Chimeras with high ES cell contribution as evidenced by coat colour were backcrossed to C57BL/6N wildtype (WT) or C57BL/6N-Tyr<c> mice for germline transmission of the targeted allele.

**Fluorescent *In Situ* Hybridization**

The dual color fluorescent *in situ* hybridization was carried out as previously described [7]. Genomic fragments containing the *nongouti* (*a*) (BAC clone B6Ng01-189F19) and *Rln3* (BAC clone B6Ng01-342B04) regions were cloned into the Charomid vector (NIPPON GENE CO., LTD., Toyama, Japan). The BAC clones were obtained from the Gene Engineering Division, RIKEN BRC. The Charomid DNA and *nonagouti* or *Rln3* BAC clones were labelled with digoxigenin-11-dUTP (Roche) or DNP-11-dUTP (Perkin Elmer) using nick translation, respectively. Anti-Dinitrophenyl (DNP) (Sigma) and Alexa Fluor 488 donkey anti-goat IgG (Invitrogen) were diluted to 1:200 and used to detect DNP-labeled *Rln3* probes. Monoclonal anti-Digoxigenin (Sigma) and Cy3-conjugated anti-mouse IgG (Jackson ImmunoResearch Inc.) were diluted to 1:300 and used to detect DIG-labeled *nonagouti* probe.

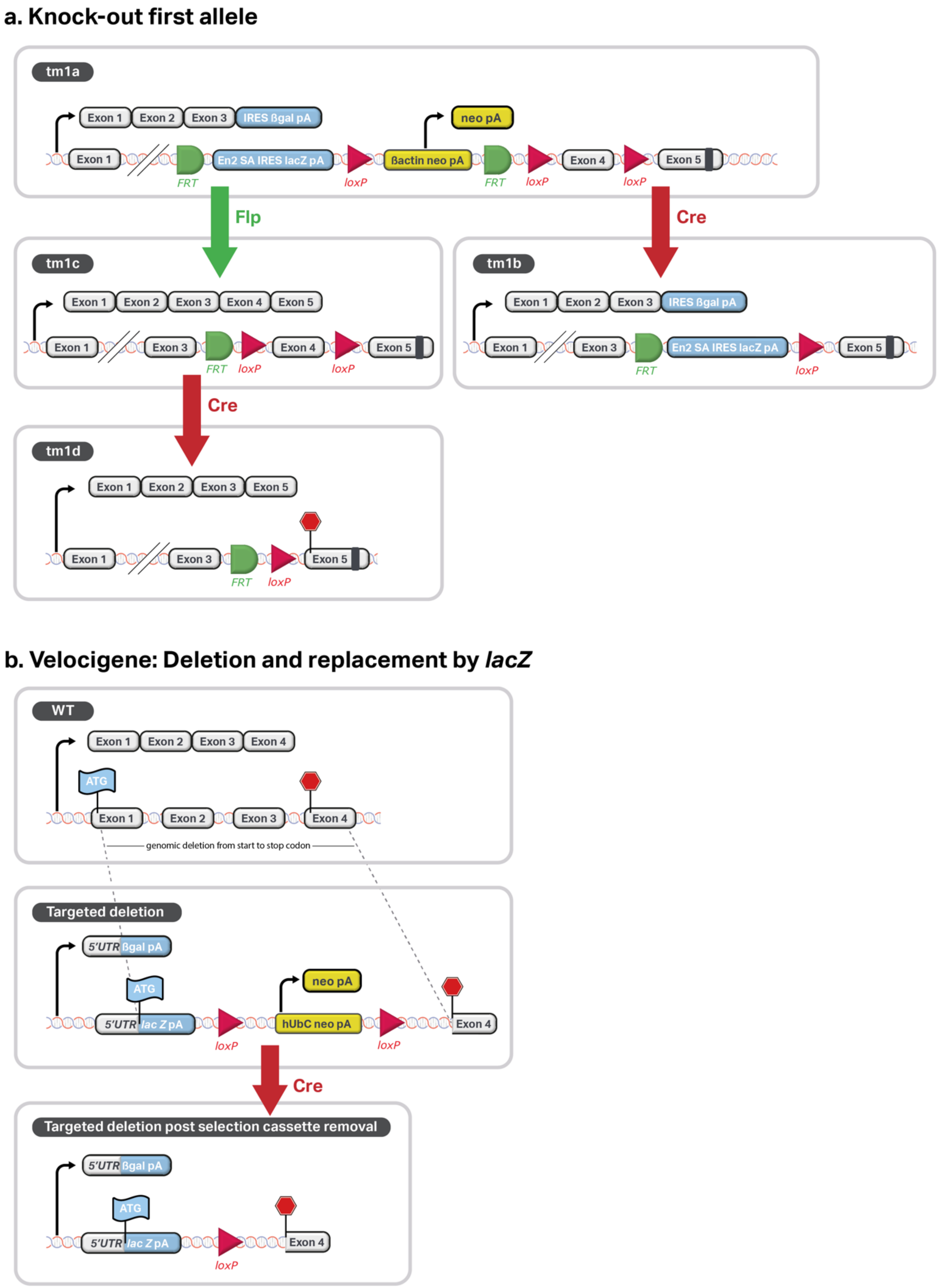


**Supplemental Figure S1. Allele types and gene products.** (a) KO-first allele and derivatives. Structure of alleles and transcripts are shown. The "KO-first" tm1a allele contains a targeted *lacZ* gene trap cassette that expresses the beta-galactosidase enzyme, a *neo* selection cassette that was used to isolate the ES cell clone, and loxP sites flanking the critical region (CR, exon(s) that will create a frameshift or eliminate an essential protein domain of the coding sequence when deleted) [9]. Cre recombinase converts tm1a alleles into a *lacZ*-tagged null tm1b allele by deleting the selection cassette if flanked by loxPs and the CR. Functional gene expression is inhibited by the trapping cassette while the risk of alternatively spliced rescue of wildtype expression is further mitigated by the deletion of the CR. The deletion also removes the strong heterologous promoter included to express the selectable marker, which can influence the expression of nearby genes [10,11]. In addition, Flp Recombinase Recognition Target (FRT) sites flank both trapping and selection cassettes, which allows for conversion of conditional-ready tm1a alleles into conditional tm1c alleles via Flp recombinase. A second recombination (by Cre recombinase) converts tm1c alleles into null tm1d alleles by deletion of the CR. Some tm1a designs employ a ß-geo coding sequence as part of the gene trap cassette and do not contain a separate selection cassette. (b) Velocigene (Vlcg) allele. The entire genomic interval that contains coding sequences is replaced by a *lacZ* cassette inserted at the ATG site of the longest transcript [2]. A selection cassette flanked by loxP sites is inserted downstream of the *lacZ* coding cassette. Cre recombinase deletes the selection cassette and strong heterologous promoter from the allele.

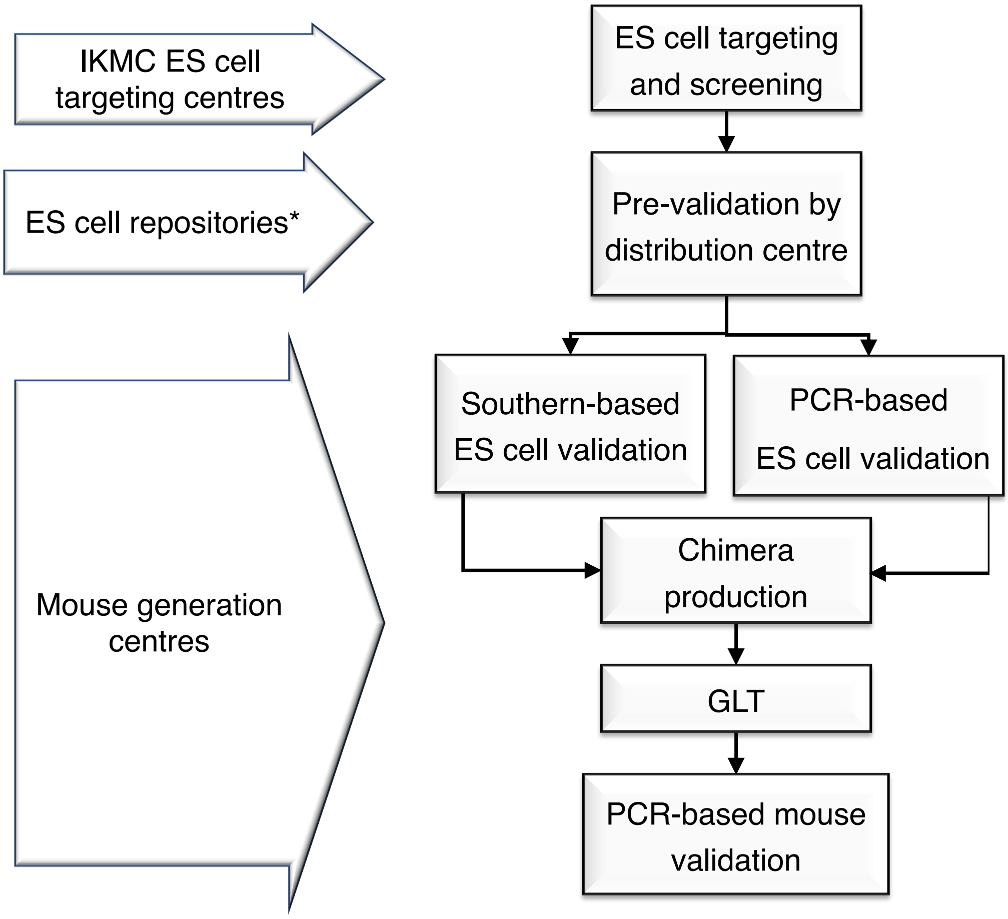


**Supplemental Figure S2**: **Process of ES cell to mouse conversion and allele validation.** Mouse productions centres received materials directly from ES cell production cores or, after a partial quality validation, from distribution hubs (Repositories). The KO-first alleles were validated by either a PCR-based or Southern-based strategy prior to chimera generation at mouse production centres. Whenever operationally feasible, ES cell chromosome count was also evaluated prior to chimera generation.

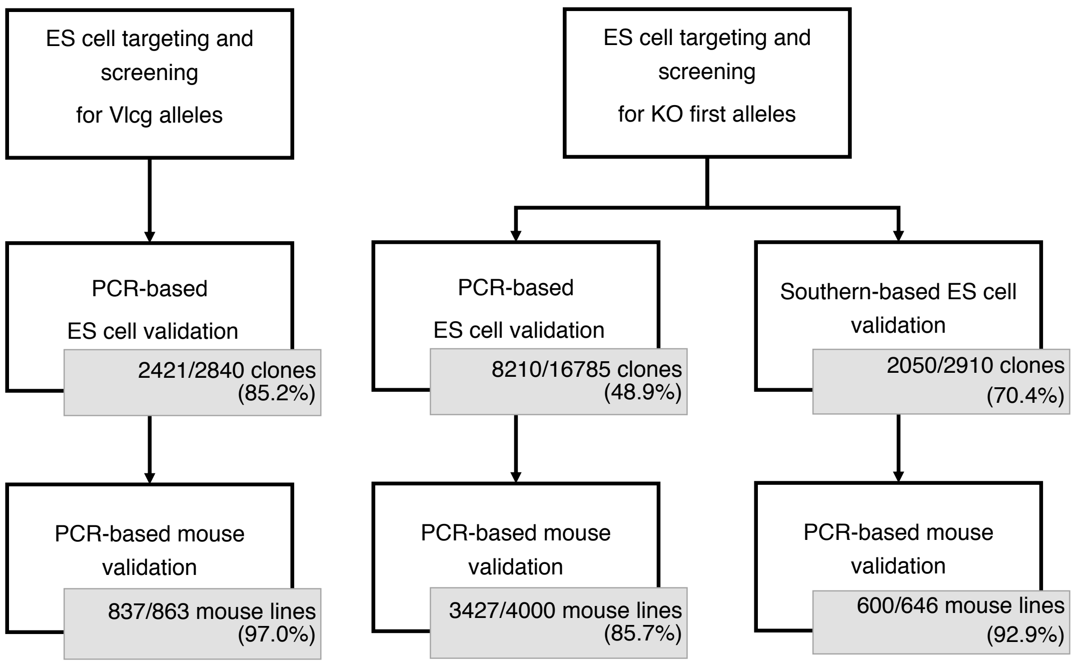


**Supplemental Figure S3: Outcome of PCR-based and Southern blot-based KO-first allele validation in ES cell clones and in mouse lines.** Validation of Vlcg deletion alleles and KO-first alleles employed distinct workflows reflecting the difference in their structure and targeting strategy. Due to the use of long homology arms (~50kb) in the targeting construct, Vlcg deletion alleles were validated by copy-counting of both non-targeted endogenous sequence (loss of allele, LOA) [2] and of the targeted cassette insertion (*lacZ*/*neo*) [8]. 85.2% of Vlcg ES clones passed secondary (pre-injection) QC, resulting in mouse lines that pass QC at a 97% overall pass rate. KO-first alleles were validated using a suite of PCR-based or Southern-based protocols, resulting in QC pass rates of 48.9% and 70.4%, respectively. Some of the difference in failure rate can be attributed to the use of pre-validated clones from a Repository. Irrespective of screening method, the significant clone rejection rate highlights the value of pre-screening ES cell clones prior to investing in chimera production. Whilst PCR-based workflows are able to frequently eliminate clones with incorrect targeting events prior to chimera production (85.6% of new mouse lines created validated), the data highlight that an ES cell validation workflow that includes analysis by Southern blot is a more stringent quality screen and better predictor of validated lines (92.9% of new mouse lines validated).

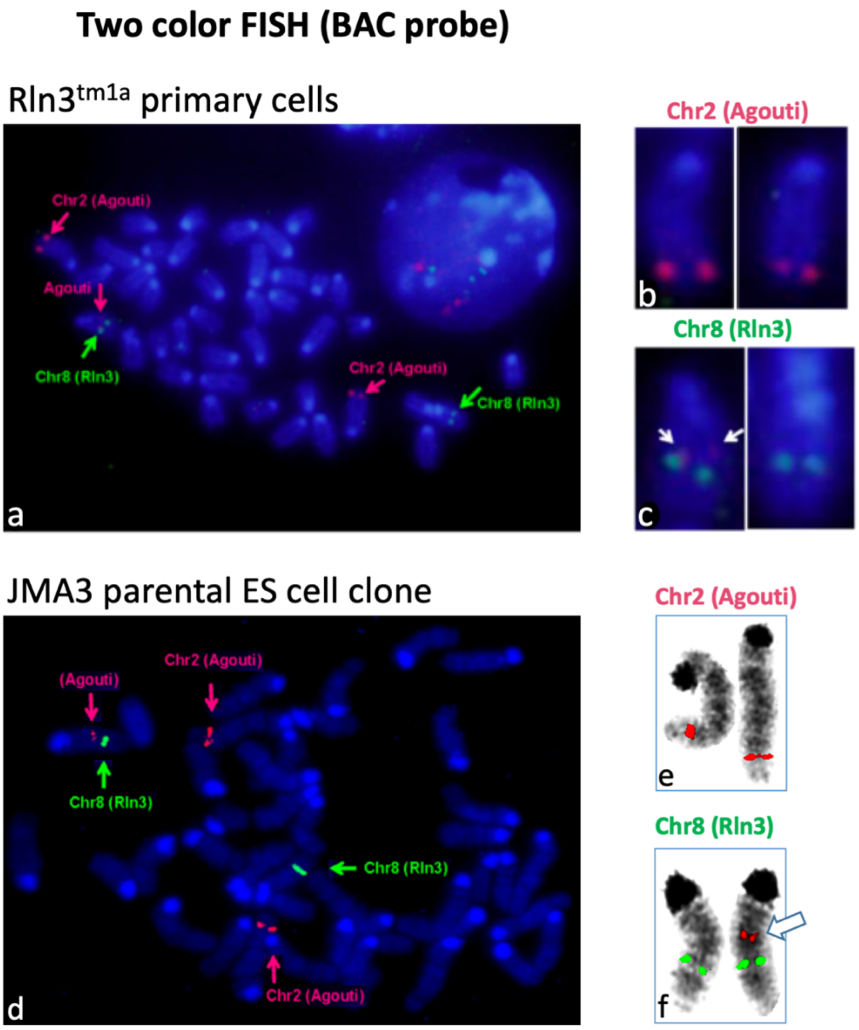


**Supplemental Figure S4: Additional *wild-type nonagouti* (*a*) insertion in JM8A3 and derivatives.** Although the entire collection was generated employing C57BL/6N ES cells, which is a nonagouti (*a*) coat colour strain, some parental clones were engineered to contain a functional *A* allele facilitating detection of germline transmission indicated by agouti coat colour[12]. Unexpectedly, some mutant lines exhibited linkage of the agouti coat colour to chromosome 8-targeted genes. Fluorescent *In Situ* Hybridization (FISH) analysis of primary cells from $Rln3^{tm1}$ animals with probes targeting the *a* gene (red) together with probes allowing the identification of chromosome 8 (green) showed copies of the *a* gene on chromosome 2 as expected (panels a and b) but also revealed additional, hemizygous copies of the *a* gene on chromosome 8 (identified by an *Rln3* probe in green) (panels a and c). To confirm if this was a unique event in this targeted line or pre-existent in the parental ES cell line, the parental ES cell line was then independently tested showing the same result (d-f). White arrows show signal from the *a* probe on chromosome 8.

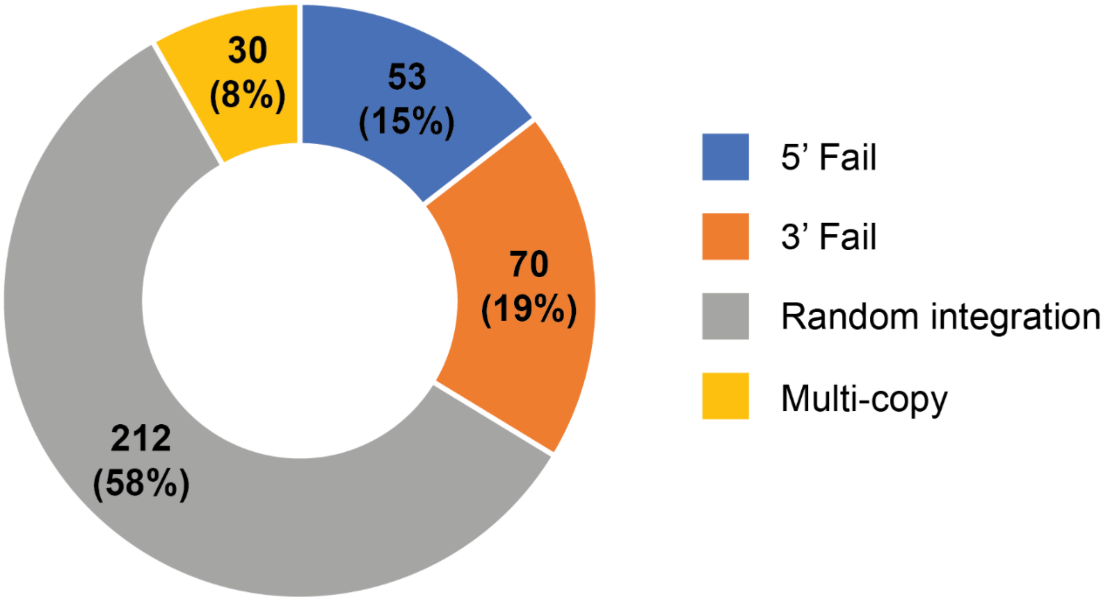


**Supplemental Figure S5: Types and percentage of observed KO-first allele failures.** Distribution of QC failure findings for 365 ES cell clones (no Repository pre-screen) analysed at a single Centre by Southern blot. The range of defects found illustrates why clone validation is a complex exercise. 5’Fail/ 3’Fail: Unexpected pattern obtained with an enzyme that cuts at sites flanking the 5’ or 3’ homology arms, respectively; Random integration: failure of both 5’ and 3’ probes. Multi-copy: evidence of multiple integrations or mixed clones.

| | JM8+subclones | | | VGB6 | | | C2 | | |
|---|---|---|---|---|---|---|---|---|---|
| Attempt number | GLT | Genes | GLT% | GLT | Genes | GLT% | GLT | Genes | GLT% |
| 1st | 2905 | 6921 | 41% | 894 | 1414 | 63% | 49 | 64 | 76% |
| 2nd | 718 | 2057 | 34% | 160 | 336 | 47% | 6 | 9 | 66% |
| 3rd | 210 | 659 | 31% | 22 | 52 | 42% | 1 | 2 | 50% |
| 4th | 71 | 261 | 27% | 6 | 14 | 42% | nd | nd | nd |
| 5 and more | 48 | 192 | 23% | nd | nd | nd | nd | nd | nd |

**Supplementary Table ST3: Germline transmission rates and numbers of clones per gene injected.** Typically, a first clone per gene was injected for chimera production and assessed for germ line transmission (GLT). In many cases, additional clones for the gene were injected if the first attempt did not yield suitable chimeras or GLT. Success rates across the Consortium, on first and subsequent attempts, indicate that injection of three to four separate clones increases the likelihood of obtaining GLT for a given gene. However, additional clones yield a diminishing return on investment in chimera production effort. Failure to obtain GLT could be for reasons intrinsic to mutant gene function or because clones for a given gene are derived from a common batch of ES cells with compromised germline potential.

| | Criterion | PCR-based strategy | Southern-based strategy |
|---|---|---|---|
| **ES cell validation** | Target on locus | LOA-qPCR and/or LR-PCR and/or Backbone srPCR | Southern blotting |
| | Number of integrations | *neo/lacZ* qPCR | |
| | Integrity / Structure of locus | srPCR | |
| | Presence of distal *loxP* | PCR (and sequencing where relevant)* | PCR (and sequencing where relevant)* |
| **Mouse validation** | Target on locus | LOA-qPCR and/or LR-PCR and/or Backbone srPCR | LOA-qPCR, srPCR |
| | Number of integrations | *neo/lacZ* qPCR | *neo/lacZ* qPCR |
| | Integrity / Structure of locus | srPCRs | srPCRs |
| | Presence of distal *loxP* | srPCR, qPCR, and/or cre deletion | cre deletion |

*Additional sequencing was performed if primers were not specific to the *loxP* sites.

**Supplemental Table ST4: Strategy for KO-first allele validation.**
Mouse production centres elected to perform allele validation with either a PCR-based or a Southern blot-based strategy to validate materials (ES cells and/or mice) against the four classical gene targeting criteria [3]. The table summarizes these strategies. LOA-qPCR=loss-of-allele qPCR assay that copy counts wildtype (WT) alleles; LR-PCR=long range PCR, with a primer outside of the homology arm and a primer specific to the targeting construct; *neo/lacZ* qPCR=copy counting of targeting construct insertion; backbone srPCR=short range PCR to detect the presence of the backbone of a targeting construct, a sign of random integration; srPCRs=a series of amplicons were amplified from functionally essential segments of alleles (e.g. FRT and loxP sites) and were occasionaly verified by Sanger sequencing; cre deletion=*loxP* integrity was verified by assessing the ability of the allele to undergo Cre-mediated conversion; Southern blotting=genomic DNA blotting analysis employed universal probes against *neo* or *lacZ* that produced a profile of bands of sizes specific to targeted allele identity. The order in which the assays were performed was different between production centres, generally starting with cost-efficient assays that did not require gene-specific reagents and performing additional tests only on materials that passed previous tests. Irrespective of assay choices, validation was essential at both the ES cell and mouse.

### Supplemental References


1. Bult, C.J. *et al.* Mouse Genome Database (MGD) 2019. *Nucleic Acids Res* **47**, D801-D806 (2019).
2. Valenzuela, D.M. *et al.* High-throughput engineering of the mouse genome coupled with high-resolution expression analysis. *Nat Biotechnol* **21**, 652-9 (2003).
3. Ryder, E. *et al.* Molecular characterization of mutant mouse strains generated from the EUCOMM/KOMP-CSD ES cell resource. *Mamm Genome* **24**, 286-94 (2013).
4. Codner, G.F. *et al.* Aneuploidy screening of embryonic stem cell clones by metaphase karyotyping and droplet digital polymerase chain reaction. *BMC Cell Biol* **17**, 30 (2016).
5. Behringer, R. *Manipulating the mouse embryo : a laboratory manual*, xxii, 814 pages (Cold Spring Harbor Laboratory Press, Cold Spring Harbor, New York, 2014).
6. Gertsenstein, M. *et al.* Efficient generation of germ line transmitting chimeras from C57BL/6N ES cells by aggregation with outbred host embryos. *PLoS One* **5**, e11260 (2010).
7. Amano, T. *et al.* Chromosomal dynamics at the Shh locus: limb bud-specific differential regulation of competence and active transcription. *Dev Cell* **16**, 47-57 (2009).
8. Tesson, L., Heslan, J.M., Menoret, S. & Anegon, I. Rapid and accurate determination of zygosity in transgenic animals by real-time quantitative PCR. *Transgenic Res* **11**, 43-8 (2002).
9. Skarnes, W.C. *et al.* A conditional knockout resource for the genome-wide study of mouse gene function. *Nature* **474**, 337-42 (2011).
10. Maguire, S. *et al.* Targeting of Slc25a21 is associated with orofacial defects and otitis media due to disrupted expression of a neighbouring gene. *PLoS One* **9**, e91807 (2014).
11. West, D.B. *et al.* Transcriptome Analysis of Targeted Mouse Mutations Reveals the Topography of Local Changes in Gene Expression. *PLoS Genet* **12**, e1005691 (2016).
12. Pettitt, S.J. *et al.* Agouti C57BL/6N embryonic stem cells for mouse genetic resources. *Nat Methods* **6**, 493-5 (2009).

**Animal Welfare Assurances**

| Institute | Information |
|---|---|
| BCM Baylor College of Medicine | Approval committee: Institutional Animal Care and Usage Committee. Approval License: AN-5896 & AN-2803 |
| GMC Helmholtz Zentrum München | Approval committee: Regierung von Oberbayern. Approval License: 144-10 |
| PHENOMIN | Approval Committee: Com'Eth N°17 and French Ministry for Superior Education and Research (MESR). Approval licenses: internal numbers 2012-009 & 2014-024. Approval licenses: MESR: APAFIS#4789-2016040511578546 |
| MRC Harwell | Approval committee: Animal Welfare and Ethical review Board (AWERB). Approval License: 30/3384 |
| Nanjing University | Approval committee: IACUC of MARC. Approval License: NRCMM9 |
| RBRC RIKEN BioResource Research Center | Approval committee: The Institutional Animal Care and Use Committee of RIKEN Tsukuba Branch. Approval License: Exp11-002, 12-002, 13-002, 14-002, 15-002, 16-002 Collection, maintenance, storage, breeding and distribution of the mouse resources Exp11-011, 12-011, 13-011, 14-009, 14-017, 15-009, 16-008 Phenotyping analyses and related studies in mice |
| The Centre for Phenogenomics | Approval committee: Animal Care Committee (ACC) of The Centre for Phenogenomics. Approval License: Animal Use Protocol (AUP) 0153, 0275, 0277, 0279 |
| The Czech Centre for Phenogenomics, Institute of Molecular Genetics of the Czech Academy of Sciences | Approval committee: IMG Committee for animal welfare and protection. Approval License: 31255/2019-MZE-18134, 16OZ9707/2019-18134 – valid till 2. 7. 2024 |
| The Jackson Laboratory | Approval: The Jackson Laboratory Institutional Animal Care and Use Committee (IACUC). License: NIH Office of Laboratory Animal Welfare (OLAW) assurance # D16-00170 IACUC Protocol: 99066<br>Accreditation: AAALACi #000096 |
| UCD University of California, Davis | Approval committee: UC Davis Institutional Animal Care and Use Committee (IACUC). Approval License: Protocol #19075 |
| WTSI Wellcome Trust Sanger Institute | Approval committee: Animal Welfare and Ethical review Board (AWERB). Approval License: PPL 80/2076 Valid 27th Nov 2006 - 3rd Jan 2012; PPL 80/2485 valid 3rd Jan 2012 - 5th Dec 2016 |
| CAM-SU | Approval committee: IACUC of CAM-SU GRC. Approval License: CAM-SU-AP#:TP-1 |
| National Laboratory Animal Center, NARLabs | Approval committee: NLAC IACUC. Approval License: 2016O05R02<br>Accreditation: AAALACi: #001204 |
| Institutes that engineer and breed mice are guided by their own ethical review panels and licensing and accrediting bodies, reflecting the national legislation under which they operate. Details of their ethical review bodies and licenses are provided here. | |

**IMPC Consortium Contributors**

*Baylor College of Medicine, One Baylor Plaza, Houston, TX, 77030, USA:* Juan J Gallegos, Jennie R Green, Ritu Bohat, Katie Zimmel

*The Centre for Phenogenomics, 25 Orde Street, Toronto, Ontario, M5T 3H7, Canada:* Monica Pereira, Suzanne MacMaster, Sandra Tondat, Linda Wei, Tracy Carroll, Jorge Cabezas, Qing Fan-Lan, Elsa Jacob, Amie Creighton, Patricia Castellanos-Penton, Ozge Danisment, Shannon Clarke, Joanna Joeng, Deborah Kelly, Christine To, Rebekah van Bruggen

*German Mouse Clinic, Institute of Experimental Genetics, Helmholtz Zentrum München, German Research Center for Environmental Health GmbH, Ingolstaedter Landstrasse 1, 85764, Neuherberg, Germany:* Valerie Gailus-Durner, Helmut Fuchs, Susan Marschall, Stefanie Dunst, Markus Romberger, Bernhard Rey

*INFRAFRONTIER GmbH:* Sabine Fessele, Philipp Gormanns

*Institute of Developmental Genetics, Helmholtz Zentrum München, German Research Center for Environmental Health GmbH, Neuherberg, Germany; Chair of Developmental Genetics, Center of Life and Food Sciences Weihenstephan, Technische Universität München, Freising-Weihenstephan, Germany:* Roland Friedel, Cornelia Kaloff, Andreas Hörlein, Sandy Teichmann, Adriane Tasdemir, Heidi Krause, Dorota German, Anne Könitzer, Sarah Weber, Joachim Beig

*The Jackson Laboratory, 600 Main Street, Bar Harbor, Maine 04609, USA:* Matthew McKay, Richard Bedigian, Stephanie Dion, Peter Kutny, Jennifer Kelmenson, Emily Perry, Dong Nguyen-Bresinsky, Audrie Seluke, Timothy Leach, Sara Perkins, Amanda Slater, Michaela Petit, Rachel Urban, Susan Kales, Michael DaCosta, Michael McFarland, Rick Palazola, Kevin A. Peterson, Karen Svenson, Robert E. Braun, Robert Taft

*Mouse Biology Program, University of California, Davis, California, 95618, USA:* Mark Rhue, Jose Garay, Dave Clary, Renee Araiza, Kristin Grimsrud, Lynette Bower, Nicole L Anchell, Kayla M Jager, Diana L Young, Phuong T Dao

*MRC Harwell Institute, The Mary Lyon Centre, Harwell Campus, Oxfordshire, OX11 0RD, UK:* Wendy Gardiner, Toni Bell, Janet Kenyon, Michelle E Stewart, Denise Lynch, Jorik Loeffler, Adam Caulder, Rosie Hillier

*MRC Harwell Institute, Mammalian Genetics Unit, Harwell Campus, Oxfordshire, OX11 0RD, UK:* Mohamed M Quwailid, Rumana Zaman, Luis Santos

*RIKEN BioResource Research Center, 3-1-1 Koyadai, Tsukuba, Ibaraki 305-0074, Japan:* Yuichi Obata, Mizuho Iwama, Hatsumi Nakata, Tomomi Hashimoto, Masayo Kadota, Hiroshi Masuya, Nobuhiko Tanaka, Ikuo Miura, Ikuko Yamada, Tamio Furuse

*Université de Strasbourg, CNRS, INSERM, IGBMC, PHENOMIN-ICS, 1, rue Laurent Fries, Illkirch, 67400, France:* Mohammed Selloum, Sylvie Jacquot, Abdel Ayadi, Dalila Ali-Hadji, Philippe Charles, Elise Le Marchand, Amal El Amri, Christelle Kujath, Jean-Victor

Fougerolle, Peggy Mellul, Sandrine Legeay, Laurent Vasseur, Anne-Isabelle Moro, Romain Lorentz, Laurence Schaeffer, Dominique Dreyer, Valérie Erbs, Benjamin Eisenmann, Giovanni Rossi, Laurence Luppi, Annelyse Mertz, Amélie Jeanblanc

*Wellcome Sanger Institute, Genome Campus, Hinxton, CB10 1SA, UK:* Evelyn Grau, Caroline Sinclair, Ellen Brown, Helen Kundi, Alla Madich, Mike Woods, Laila Pearson, Danielle Mayhew, Nicola Griggs, Richard Houghton, James Bussell, Catherine Ingle, Sara Valentini, Diane Gleeson, Debarati Sethi, Tanya Bayzetinova, Jonathan Burvill, Bishoy Habib, Lauren Weavers, Ryea Maswood, Evelina Miklejewska, Ross Cook, Radka Platte, Stacey Price, Sapna Vyas, Adam Collinson, Matt Hardy, Priya Dalvi, Vivek Iyer, Tony West, Mark Thomas, Alejandro Mujica, Elodie Sins,Daniel Barrett

*Phenomics Australia, John Curtin School of Medical Research, The Australian National University, 131 Garran Road, Acton, Canberra, ACT 2601, Australia:* Michael Dobbie

*PCDDP, North-West University, 11 Hoffman street, Potchefstroom, 2531, South Africa:* Anne Grobler, Glaudina Loots, Rose Hayeshi, Liezl-Marie Scholtz, Cor Bester, Wihan Pheiffer, Kobus Venter

*Universitat Autònoma de Barcelona:* Fatima Bosch
